# Chronic compression drives macrophages toward a pathological pro-tumor state

**DOI:** 10.1101/2025.05.13.653793

**Authors:** Alice Burchett, Hao Chen, Julian Najera, Scott Howard, Meenal Datta

**Affiliations:** Department of Aerospace and Mechanical Engineering, University of Notre Dame, IN, USA; Department of Electrical Engineering, University of Notre Dame, IN, USA

**Keywords:** Myeloid cells, solid stress, immunomechanics, mechano-immunology, metabolism, fluorescence lifetime imaging (FLIM), phagocytosis, polarization, RNA sequencing, morphology

## Abstract

Macrophages comprise a significant portion of the glioblastoma tumor microenvironment and are essential in promoting immunosuppression and tumor progression. Solid tumors such as glioblastoma generate solid stress as they expand, creating a compressive microenvironment for mechanosensitive immune cells including macrophages. Macrophages are known to respond to various mechanical stimuli but have not yet been studied in the context of chronic compression observed in growing tumors. Here, we used a custom *in vitro* compression system to elucidate the effects of compressive solid stress on murine macrophages. We found that macrophages have significant morphological, transcriptional, metabolic, and functional responses to compression. These changes corresponded to both canonical pro- and anti-inflammatory macrophage states. The gene expression signatures of compressed macrophages more closely resembled those of glioma-associated macrophages known to be associated with worse patient outcomes. These results indicate that compression alone, independent from tumor cell-derived biochemical factors, may contribute to the pathological tumor-associated macrophage phenotype. This could represent a vicious cycle of tumor immunomechanics and mechano-immunology. Targeting solid stress in tumors or the response to solid stress by macrophages may interrupt this feedback loop to help normalize the tumor immune microenvironment and improve glioblastoma response to immunotherapy.

## Introduction

Myeloid cells, such as macrophages, accumulate in the glioblastoma (GBM) tumor microenvironment (TME) and can comprise nearly half of the tumor bulk (1,2). Macrophage phenotype is often represented as ranging from pro-inflammatory (M1-like) to anti-inflammatory (M2-like), with intermediate phenotypes found in both health and disease (3). In GBM, they adopt a pathological, M2-like pro-tumor phenotype (2). As key orchestrators of the innate and adaptive immune response, these tumor-associated macrophages (TAMs) inhibit an effective anti-tumor immune response and contribute to tumor growth, invasion, and treatment resistance (4). They are also key players in resistance to immunotherapies (5). As a result, TAM density is associated with poor prognosis and reduced overall survival in many cancer types (4,6,7).

The GBM TME also features elevated solid stress, which arises as the tumor expands and pushes against the surrounding tissue (8–10). This chronic compression not only acts on the surrounding brain to cause neuronal death and inhibit fluid transport (11,12), but also acts on the TME-resident macrophages. Macrophages are known to respond to mechanical stimuli, with the polarization and phenotype response highly dependent on the physiological context (13,14). We recently showed that macrophages can themselves generate solid stress *de novo*, causing deformation of a confining gel depending on gel stiffness and biochemical cues (15). While several other studies have explored the macrophage response to chronic compression in the context of bone and tooth remodeling, there are no studies to date that define their response to compressive stress in the context of cancer immunology (16–19).

Macrophage function is closely linked to metabolic state. M2-like polarization is accompanied by an increase in oxidative phosphorylation compared to more glycolytic M1-like macrophages (20–22). While TAMs are often observed to rely on glycolysis for energy production, TAMs in hypoxic environments and in the core of GBM tumors have increased oxidative phosphorylation (23–25). Macrophages and GBM cells participate in reciprocal metabolic cross-talk, resulting in a vicious cycle that promotes GBM growth and invasiveness and accumulation of pro-tumor, immunosuppressive TAMs (23). The mechanical properties of the TME are also directly interconnected with tumor cell metabolism (26). Macrophages within stiffened, fibrotic TMEs upregulate arginase-1 and metabolize arginine, leading to the depletion of arginine in the TME (27). This suppresses anti-tumor T cell activity and leads to the production of pro-tumor metabolites. Because macrophages can themselves generate compressive solid stress (15), there may be an immunomechanical and metabolic feedback loop involving the GBM mechanical TME and tumor-resident macrophages resulting in immunosuppression and tumor progression.

Here, we utilize a 2D mechanical compression model to interrogate macrophage response to chronic solid stress levels previously measured in GBM. Compression induced alterations in macrophage morphology, including increased cell area and more irregular cell shapes. Compressed macrophages also had significantly altered gene expression as assessed by RNA sequencing, upregulating both M1-like and M2-like genes, defying the simplified M1/M2 macrophage polarization spectrum. They also upregulate genes associated with TAMs across a spectrum of tumor types, including glioma and GBM-specific TAMs. Trending upregulation of canonical M1-like and M2-like markers was consistent across varying magnitudes of solid stress, duration of compression, and in normoxic and hypoxic conditions. Fluorescence lifetime imaging (FLIM) showed alterations in cellular metabolism under compression, as did transcriptional analysis. Functionally, compressed macrophages were more phagocytic and released more nitric oxide into their media. These results suggest that chronic compression results in a more complex phenotype than is represented on the traditional pro- and anti-inflammatory axis. This also demonstrates that compression alone, in the absence of any cancer-derived factors, can cause macrophages to adopt a TAM-like state, implicating solid stress as a potentially essential driver of tumor immunosuppression and a promising therapeutic target.

## Materials and Methods

### Cell culture and compression

RAW264.7 murine macrophages were purchased from ATCC (TIB-71). Cells were cultured in Dulbecco’s Modified Eagle Medium (DMEM, Corning, 31053-028) supplemented with fetal bovine serum (FBS, Corning, 35-010-CV) and penicillin/streptomycin (Corning, 30-002-cl). Unless otherwise noted, cells were maintained at 37°C and 5% CO_2_ in a humidified incubator.

We employed a custom weight system to apply uniaxial compressive stress to cell monolayers. Cells were seeded atop a cell culture insert with 0.4 um pore size (CellQuart, 9300412) and placed in a 6-well plate. Agarose cushions were made from 1% (w/v) agarose (Bio-Rad, 1613100) dissolved in complete culture medium and formed with custom 3-D printed cylindrical molds to match the size of the cell culture insert. These serve as a media reservoir to prevent nutrient deprivation and protect the cells from direct contact with the rigid weight. The agarose cushions were submerged in culture medium and placed in the incubator for 24 hours before use to equilibrate CO_2_ and oxygen within the gels. After cells had adhered to the cell culture insert (overnight), the agarose cushion was placed on top, and a custom 3-D printed PLA weight was placed on top of the agarose. The mass of the weight is such that it applies 0.14 kPa of compressive stress distributed across the insert membrane area. For the control condition, only the agarose cushion was placed on the cells. After the addition of the agarose (control) or agarose + weight (compressed), the cells were placed in a standard incubator for 24 or 48 hours. For the hypoxia condition, the cells were placed in a hypoxia incubator set at 1% O_2_ and 5% CO_2_. Conditioned media was collected from the well underneath the cell culture insert and stored at −80°C before downstream analysis. For M1-polarized macrophages, media was supplemented with 200ng/ml lipopolysaccharide (LPS, Santa Cruz Biotechnology, sc-3535) and 20 ng/ml IFN-γ (BioLegend, 575302). M2-polarized macrophages were treated with 20 ng/ml IL-4 (Pepro-tech, 214-14).

### Cell morphology analysis

After compression, the cells were fixed with 4% paraformaldehyde (Thermo Scientific, AAJ19943K2) for 20 minutes at room temperature and rinsed with PBS. The membranes were removed from the cell culture inserts and stored in PBS with 0.1% sodium azide (Sigma-Aldrich, S2002). Phase-contrast images of these fixed membranes were acquired on a Leica DMi 1 microscope. The membranes were cut into several pieces using a razor. Single pieces of the membranes were placed into a 12-well plate for staining using conjugated antibodies against YAP (Santa Cruz Biotechnology, sc-376830), TAZ (Santa Cruz Biotechnology, sc-518026), and DAPI (Sigma-Aldrich, D9542). The stained membranes were mounted onto microscope slides under coverslips using Fluoromount G (Invitrogen, 00-4958-02). Image Z-stacks were acquired using a Nikon AXR confocal microscope, and the maximum intensity projections were used for downstream analysis. The YAP, TAZ, and DAPI channels were merged to obtain images of the entire cell, and images were cropped to regions that did not contain overlapping cells for more accurate segmentation. The cropped images were segmented using CellProfiler, which quantified cell area and shape metrics. Representative images shown were created by staining cells with CellMask actin tracking stain (Invitrogen, A57244) and DAPI.

### Fluorescence lifetime imaging

For fluorescence lifetime imaging (FLIM), cells were seeded sparsely in a 12-well cell culture insert and compressed as described above. After 24 or 48 hours, the weight and agarose cushion were carefully removed and the cells were fixed as described above. The membrane was removed from the cell culture insert and mounted using Fluoromount G onto glass microscope slides. The coverslip edges were sealed with nail lacquer to prevent moisture from reaching the membranes. The slides were stored at 4°C before imaging.

FLIM measures the inherent lifetime (τ) of fluorophores regardless of intensity, enabling tracking of bio-microenvironment dynamics. In this study, we utilized a custom-built two-photon excited frequency-domain FLIM system. “Instant-FLIM” captures intensity and lifetime imaging simultaneously under 800 nm excitation for intrinsic fluorescence lifetime through precise measurements of endogenous fluorophores such as NAD(P)H (28). For each measurement, lifetime phasor coordinates g and s [2] are averaged separately to calculate the mean lifetime per field of view (FOV) for quantitative analysis (29).

### Phagocytosis and Griess assay

Cells were either compressed as described above or treated with conditioned media for 24 hours. After treatment, cells were incubated with 1 µm fluorescent polystyrene beads (Invitrogen, F13083) for 2 hours. The cells were then fixed, stained with CellBrite blue (Biotium, 30024), and imaged. We used CellProfiler to quantify the intensity of red fluorescence within the blue cell area. The conditioned media of compressed macrophages was analyzed using a Griess reagent system following the vendor protocol (Promega, G2930).

### RNA sequencing and bioinformatics analysis

RNA was extracted as described above from six samples each of 24h compressed (0.14 kPa) and 24h agarose-only control macrophages and sequenced using the NovaSeq X Plus 10B flow cell at a depth of approximately 50 million average raw reads per sample. The raw data was aligned to the GENCODE mouse genome annotation using STAR 2.7.2. The resulting expression data was analyzed using DESeq2 to identify differentially expressed genes (DEGs), create the volcano plot, create heatmaps, perform gene ontology and geneset enrichment analyses, and generate log_2_(fold change) values for specific genes. For geneset analysis, genesets were either downloaded from the GSEA online database or obtained from relevant publications (Supplementary Table 1). The protein-protein interaction network was created using STRING-db, filtering for the subset of DEGs with an absolute value of log_2_(fold change) greater than 2 and an adjusted p-value (padj.) less than 0.05. The interactions were further filtered to include only genes with a high confidence interaction score (greater than 0.9) and disconnected nodes were excluded. Violin plots were created using GraphPad Prism (GraphPad Software, San Diego, CA, USA).

### Reverse-transcription quantitative polymerase chain reaction

After completion of the compression experiment, the agarose cushion and weight were removed. The cells were removed from the membrane of the cell culture insert using a cell scraper. These were pelleted, and the RNA was extracted (Zymo Research, R2051). Gene expression was analyzed using TaqMan primers for Arg1 (Thermo Fisher Scientific, Mm00475988_m1), Nos2 (Mm00440502_m1), and GAPDH (Mm99999915_g1). The raw PCR data was analyzed using the qPCR Design and Analysis app (Thermo Fisher Scientific). Gene expression was normalized to GAPDH and then normalized to the mean of the control group and reported as 2^− ΔΔCt^. Each condition and gene had between 3 and 6 biological replicates.

### Statistical analysis

Statistical analyses and data visualization were done using GraphPad Prism. Cell size, circularity, solidity, and fluorescence lifetime were compared using a Mann-Whitney test. Phagocytosis fluorescence intensity and nitrite in media were compared using an unpaired t-test. Error bars represent standard error of the mean and asterisks indicate statistical significance (*p ≤ 0.05, **p ≤ 0.01, ***p ≤ 0.001, ****p ≤ 0.0001). For RT-qPCR data, asterisks indicate statistical significance adjusting for multiple comparisons via multiple t-tests, with the P-value threshold of 0.0167 for variable compression and 0.025 for 48h compression and hypoxia.

## Results

### Compressed macrophages have increased size and irregular morphology

Both compressed and control macrophages maintain adherence to the surface of the transwell insert, even after removal of the weight and agarose cushion, allowing for downstream staining and morphology analysis. Murine RAW264.7 macrophages were first compressed with 0.14 kPa of compressive stress for 24 hours under standard culture conditions. This magnitude of compression approximates the magnitude we previously measured in murine glioblastoma models (11,30,31). After compression, macrophages were processed for downstream transcriptomic, imaging, and functional assays (**Figure 1A**). Quantification of fluorescent staining (**Figure 1B**) reveals that compression caused cell area to nearly double, from 87.0 µm^2^ to 168.3 µm^2^ (**Figure 1C**). This is consistent with a more M1-like polarization (32). We also quantified cell circularity, defined in CellProfiler as Form Factor, or 4*π*Area/Perimeter^2^. For a perfect circle, this value equals 1, with smaller values indicating more irregular shapes. Circularity was significantly decreased in the weighted condition (**Figure 1D**). Decreased circularity has been observed with M2-like polarization (32). Solidity is the ratio of the cell area to the convex hull area (the smallest convex shape that would contain the cell) and is a metric that captures irregularity in the cell boundary, unlike size and circularity. Solidity significantly decreased in compressed macrophages compared to uncompressed controls, indicating increased cell boundary irregularity (**Figure 1E**). This is also consistent with M2-like polarization (32). Compressed macrophages had no significant change in elongation or eccentricity (data not shown).

**Figure 1.**
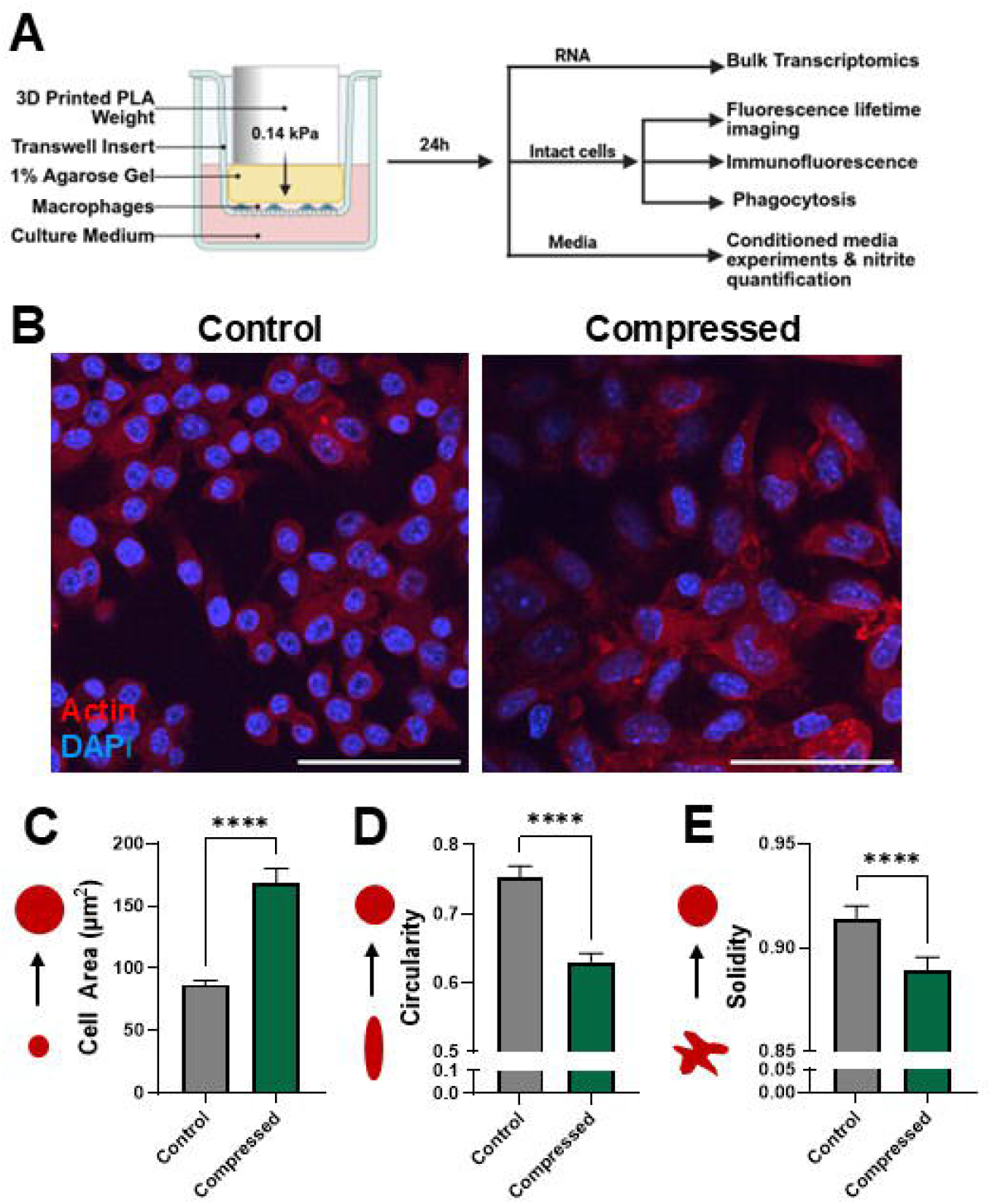
Compressed macrophages have increased size and more irregular morphology. (**A**) Schematic of the macrophage compression system and downstream analyses. Cells are seeded in a monolayer on a porous cell culture insert, and 0.14 kPa compression is added via a 3D-printed weight. The compressed macrophages are either lysed for RNA extraction, or fixed on the membrane for imaging. Conditioned media is collected for nitrite quantification and conditioned media experiments. (**B**) Fluorescence images of control and compressed macrophages stained for actin (red) and nuclei (blue). Scale bar is 50 µm. Quantification of cell area (**C**), circularity (**D**), and solidity (**E**), quantified using an automated CellProfiler pipeline which identifies and characterizes individual cells in fluorescence images (N = 94, 110 cells per group). ****p≤0.0001.

### Compression induces altered expression of genes related to migration, proliferation, and IL-1β signaling

To determine macrophages’ transcriptional response to chronic compression, we performed bulk RNA sequencing on macrophages after 24 hours of compression compared to uncompressed cells under agarose only. There were 7,463 significantly differentially expressed genes (DEGs) (p.adj < 0.05). Of these DEGs, 129 had a log_2_(Fold Change) value below −1, and 319 genes had a log_2_(Fold Change) above 1 (**Figure 2A**). The compressed and control groups were clearly delineated on a heatmap of DEGs (**Figure 2B**). We next performed gene ontology analyses to determine which relevant biological and molecular functions were altered (**Figure 2C**). The most upregulated biological processes were mainly related to migration and chemotaxis, behaviors that are more prominent in M2-like macrophages (33). Downregulated processes were mostly involved in proliferation, an effect observed in macrophages treated with pro-inflammatory stimuli (34). The molecular functions that were up- and down-regulated corroborate these results, with increased transcription of cytokine and signaling activity, and decreased DNA-related functions. To visualize the interaction between migration and proliferation, we generated a protein-protein interaction map (**Figure 2D**). Interestingly, the DEGs involved in these two processes shared many common nodes, with Il1b appearing as the most central node.

**Figure 2.**
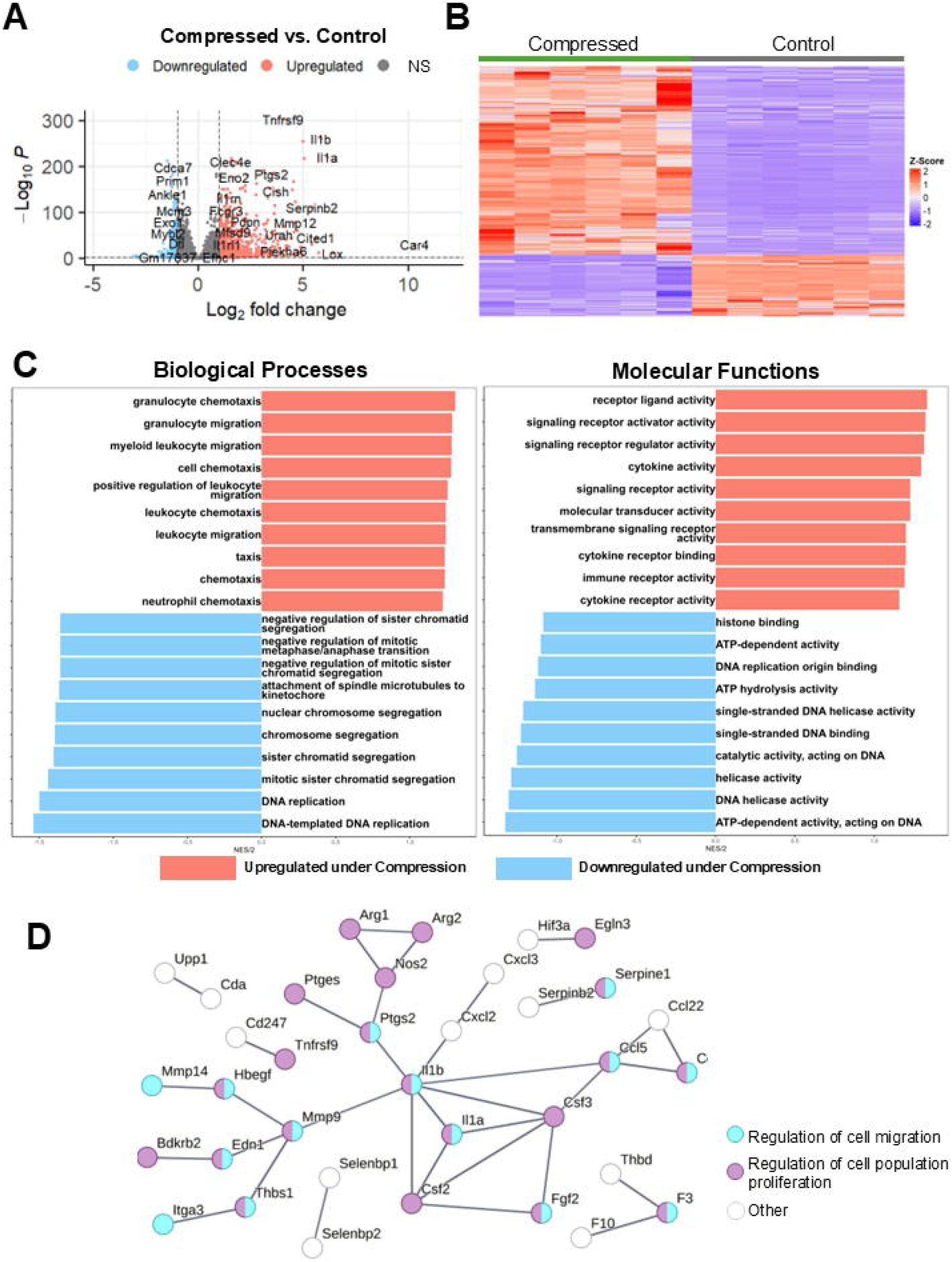
Compression upregulates gene expression related to migration and IL-1β signaling and downregulates proliferation-related genes. (**A**) Volcano plot of the top differentially expressed genes from bulk RNA sequencing of compressed versus control macrophages, with upregulated genes shown in red, downregulated genes shown in blue, and genes with padj. > 0.05 or absolute value of log_2_(fold change) < 1 shown in grey. (**B**) Heatmap showing DEGs in compressed versus control samples. (**C**) Gene ontology analysis of the top 10 up- and down-regulated (red and blue, respectively) biological processes and molecular functions in compressed versus control macrophages (**D**) Protein-protein interaction map showing DEGs with strong interactions, colored based on involvement in regulation of cell migration (cyan) and regulation of cell population (purple). N=6 per group.

### Compression induces enrichment of polarization, functional, and TAM-associated genesets

We next performed geneset variation analysis to determine changes in specific pathways of interest, beginning with macrophage polarization. Although dividing macrophage phenotype into M1-like versus M2-like is an oversimplification of the complex macrophage polarization spectrum, we compared our macrophage expression data to published M1 and M2 genesets to obtain a simplified readout of pro- or anti-inflammatory status. We chose genesets that were created as a result of a meta-analysis of murine macrophage polarization experiments, choosing a geneset that was shared between *in vitro* and *in vivo* M1 and M2 polarized macrophages (35). Compressed macrophages had significant enrichment of both M1-like and M2-like signatures compared to uncompressed controls (**Figure 3A**).

**Figure 3.**
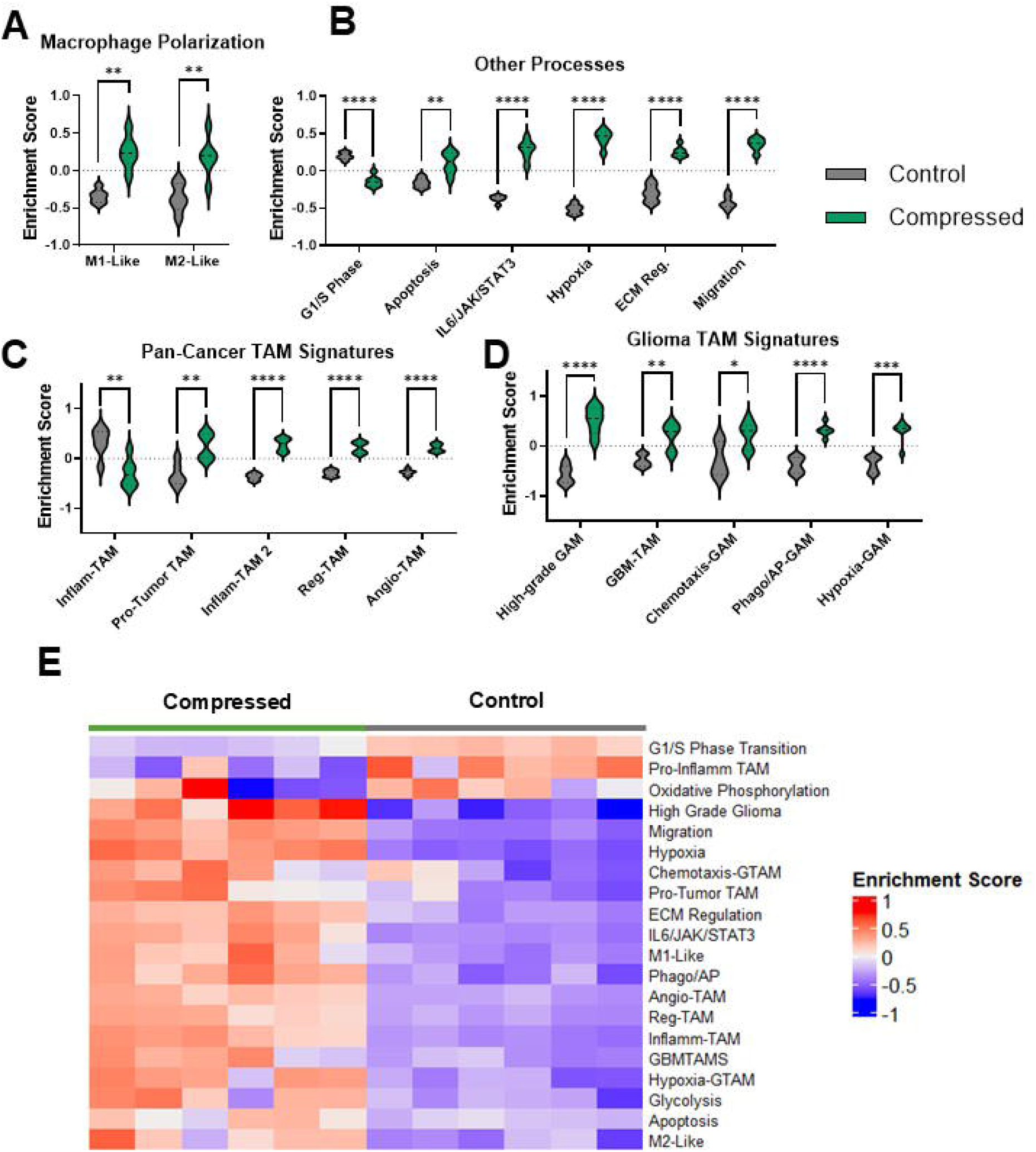
Geneset enrichment analysis reveals upregulation of polarization, functionalization, and TAM-associated genesets. Enrichment analyses of compressed versus control macrophages for M1-like and M2-like gene sets from literature (**A**), other biologically relevant pathways from the GSEA online public database (**B**), tumor-associated macrophage signatures from literature (**C**), and glioma-specific tumor associated macrophage signatures from literature (**D**). (**E**) Heatmap of enrichment scores for the gene sets shown above. Asterisks indicate statistical significance based on the p-values adjusted for multiple comparisons (N=6 per group, *p adj≤ 0.05, **padj ≤ 0.01, ***padj ≤ 0.001, ****padj ≤ 0.0001).

We next conducted analyses on pathways associated with G1-S phase transition, apoptosis, IL-6/JAK/STAT signaling, hypoxia response, extracellular matrix (ECM) regulation, and migration (**Figure 3B**). There was a significant reduction in G1/S phase enrichment with compression, suggesting reduced proliferation. G2, M2M, and general cell cycle genes were also downregulated with compression (data not shown). There was a concurrent upregulation of the apoptosis pathway. We also observed a significant increase in migration-associated genes. We found upregulation of the IL6/JAK/STAT signaling pathway, which is important in regulating macrophage inflammatory response and implicated in cancer immunity dysfunction (36). We observed an upregulation of ECM-modulating genes, also consistent with a migratory phenotype. Finally, there was an upregulation in hypoxia response genes. It is unclear whether this is in response to hypoxic oxygen levels, or mechanically-induced HIF-1α activation as observed in other mechanical settings (37–39).

Because the coinciding upregulation of both M1-associated and M2-associated genes has been observed in tumor-associated macrophages (40), we next tested our expression data against a set of tumor-associated macrophage signatures. First, we compared signatures associated with pro-inflammatory TAMs and pro-tumor TAMs, identified in a pan-cancer analysis (41). We found that the pro-inflammatory (Inflam-TAM) signature was significantly decreased in compressed macrophages, while the pro-tumor signature was significantly increased (**Figure 3C**). Notably, this pro-tumor TAM signature is associated with a worse prognosis and response to immunotherapy (41). We then obtained TAM subset gene signatures from a second pan-cancer analysis for macrophages with an inflammatory, M1-like phenotype (Inflam-TAM 2), macrophages with an immune-regulatory and M2-like phenotype (Reg-TAM), and macrophages associated with angiogenesis (Angio-TAM) (42). These three signatures were significantly upregulated with compression. Here, the upregulation of one pro-inflammatory TAM signature and the downregulation of another pro-inflammatory TAM signature derived from a separate study highlight the complexity of macrophage immune regulation and heterogeneity even in pan-cancer analyses.

We next found published genesets for glioma TAMs (GAMs), finding that compression caused a significant upregulation of several GAM subtypes (**Figure 3C**). Genes associated with macrophages found in the high-grade glioma environment (high-grade GAMs) were enriched upon compression (**Figure 3D**) (43). We also analyzed a geneset associated with macrophages in GBM-bearing mice (GBM-TAMs) compared to healthy controls, finding that it was significantly upregulated in compressed macrophages (44). Finally, we used the mouse ortholog version of genesets defining three different GAM subtypes based on analysis of several human gliomas (45). Compressed macrophages had enrichment of GAM-specific signatures relating to chemotaxis, phagocytosis/antigen presentation (Phago/AP), and hypoxia response. A heatmap of the enrichment scores for each pathway demonstrates a clear visual delineation of compressed versus uncompressed conditions (**Figure 3E**).

### Compression induces conserved polarization changes under varying magnitudes and duration of solid stress and exposure to hypoxia

We next obtained relative expression estimates of specific genes of interest from the sequencing data (**Figure 4A**). *Arg1* and *Nos2* (canonical markers of M2-like and M1-like polarization, respectively) were both significantly upregulated. *Cd274* (encoding PD-L1), *Cd47*, and *Sirpa*, all implicated in tumor immune escape (46,47), were also all significantly upregulated. However, both *Casp3* and *Mki67* were downregulated.

**Figure 4.**
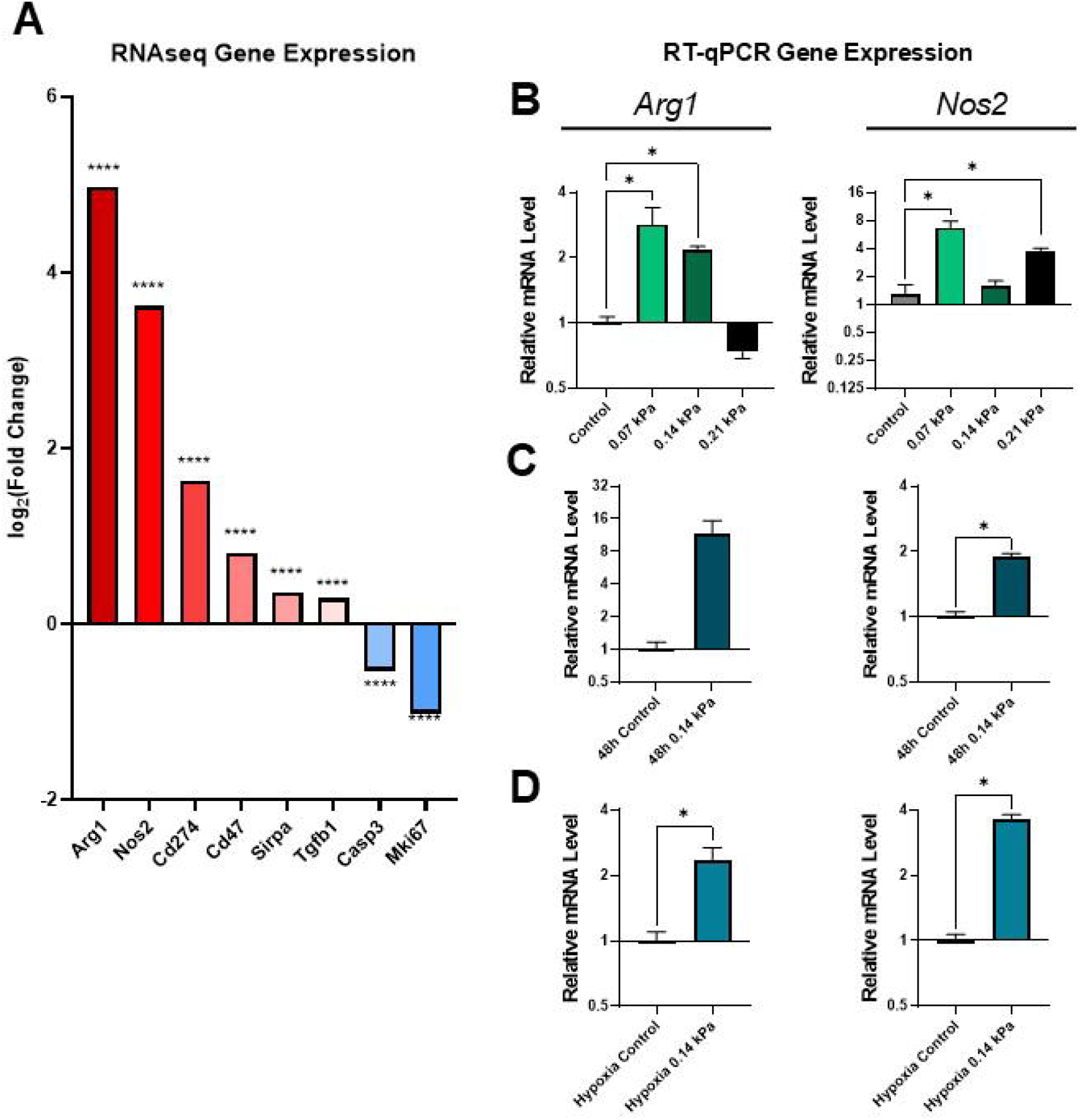
Compression upregulates both pro-/anti-inflammatory and immune evasion-related genes independent of compression magnitude duration, or oxygen levels. (**A**) Log_2_(fold change) of genes of interest based on RNA sequencing data. ****padj ≤ 0.0001, N=6. RT-qPCR quantification of *Arg1* and *Nos2* expression under various magnitudes of solid stress (**B**), increased duration of solid stress (**C**), and under simultaneous compression and hypoxia (**D**) (*p ≤ 0.05, adjusted for multiple comparisons, N=3-6 per group).

To determine whether the response to compression varies with compression magnitude, we applied 0.07, 0.14, and 0.21 kPa of compression to macrophages. There was a general trend towards increased *Arg1* and *Nos2* with compression (**Figure 4B**). Interestingly, the largest increase in *Arg1* occurred in the lowest compression group, showing a 2.9-fold increase under 0.07 kPa of compression. However, at 0.21 kPa, there was a non-significant decrease in *Arg1. Nos2* was significantly upregulated in the 0.07 kPa and 0.21 kPa conditions by 6.7 and 1.6-fold, respectively, with a non-significant slight increase with 0.014 kPa. After 48 hours of 0.14 kPa compression, *Arg1* had a large, but not statistically significant, 11.6-fold increase, while *Nos2* had a statistically significant 1.9-fold increase (**Figure 4C**). To determine whether this effect would remain under hypoxia, such as what is observed in solid tumors, we applied 0.14 kPa of compression to macrophages for 24 hours cultured in 1% O_2_ (**Figure 4D**). Both *Arg1* and *Nos2* showed statistically significant increases of 2.3-fold and 3.6-fold, respectively. While the biological variability of the macrophage transcriptional response to compression hindered the achievement of statistical significance in some conditions, these data generally demonstrate an upregulation of canonical markers of both pro- and anti-inflammatory macrophage phenotypes under various compression conditions.

### Compression induces increased average fluorescence lifetime associated with pro-tumor metabolism

NAD(P)H, a metabolic coenzyme, has an autologous fluorescence signal with variable fluorescence lifetime depending on whether it is free or bound to a protein (48). NAP(P)H in the bound state has a longer fluorescence lifetime and indicates oxidative phosphorylation, while free NAD(P)H is involved in glycolysis. Because NAD(P)H is the predominant intracellular auto-fluorescent molecule, measuring the average fluorescence lifetime of a cell indicates whether it relies more on oxidative phosphorylation (OXPHOS) or glycolysis. Macrophages have a significant shift in lifetime with polarization, where M1-like macrophages tend to have a lower lifetime and increased reliance on glycolysis while M2-like macrophages have an increased lifetime and rely on oxidative phosphorylation instead (49). We successfully generated fluorescence lifetime microscopy (FLIM) images of compressed and uncompressed macrophages (**Figure 5A**). We then obtained the average fluorescence lifetime of the cell area for each image. We observe that compressed macrophages significantly increase average fluorescence lifetime compared to uncompressed controls, a shift which is consistent with a more M2-like polarization (**Figure 5B**). We confirmed that macrophages induced to have an M2-like polarization (treated with IL-4) had a higher average lifetime than M1-like macrophages (treated with LPS and IFN-γ), consistent with prior reports (49) (**Figure 5C**).

**Figure 5.**
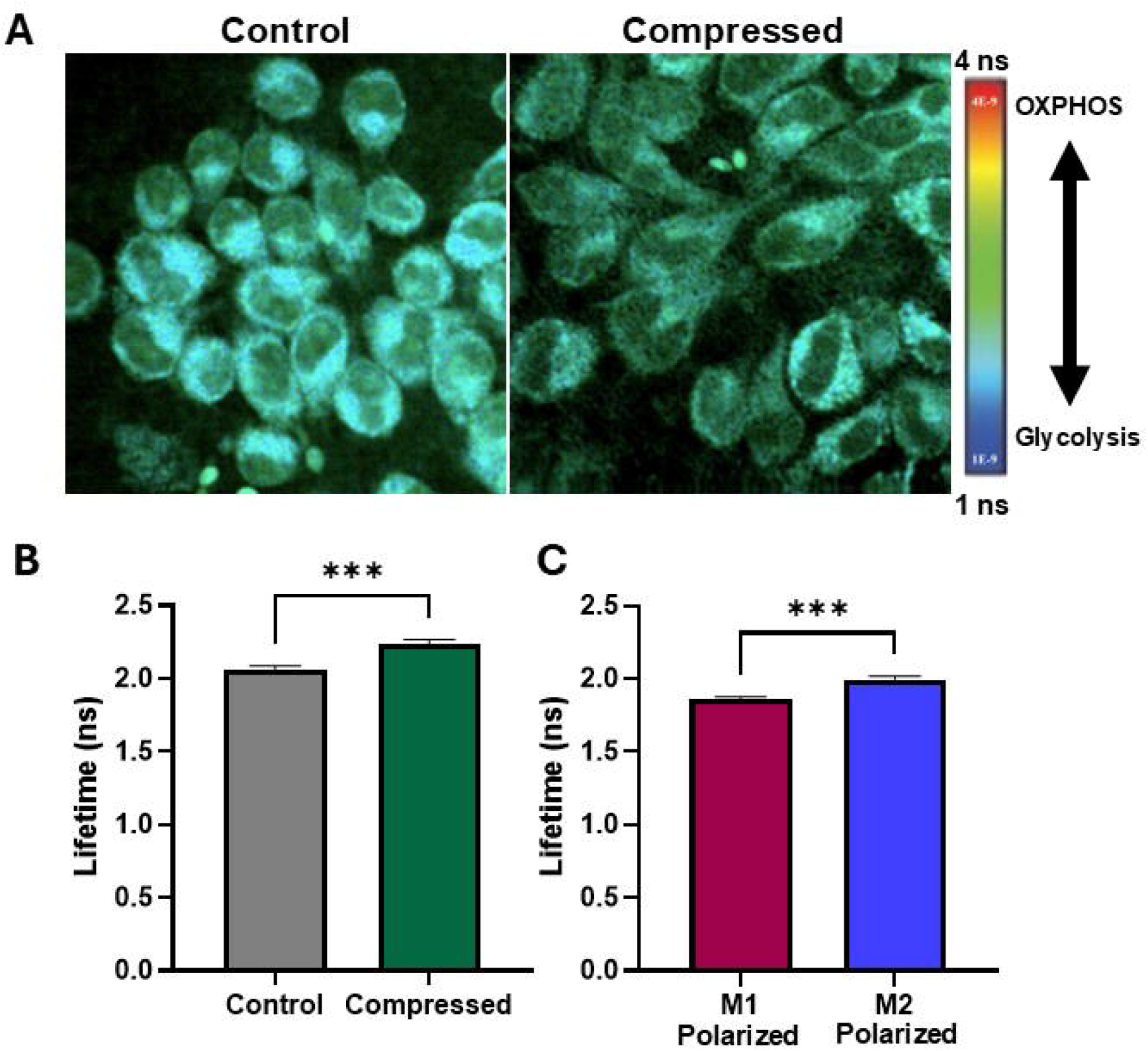
Compression increases fluorescence lifetime of macrophages, indicating increased oxidative phosphorylation and pro-tumor metabolism. (**A**) Fluorescence lifetime images of uncompressed and compressed macrophages. Cooler colors indicate lower fluorescence lifetime and increased reliance on glycolysis, and warmer colors indicate increased lifetime and increased reliance on oxidative phosphorylation (OXPHOS). (**B**) Mean fluorescence lifetime in compressed versus control macrophages. (**C**) Mean fluorescence lifetime in macrophages treated with M1 and M2 polarizing cytokines, confirming predicted upward shift with M2-polarization observed in literature (***p≤0.001, N=13-19 images per group).

### Compression promotes an increase in phagocytosis and nitric oxide production, indicative of pro-tumor function

Finally, to determine how compression alters macrophage function, we performed a phagocytosis assay on macrophages co-cultured with fluorescent microbeads immediately after 24 hours of compression. We observed a significant increase in microbead uptake in compressed macrophages (**Figure 6A, B**). Increased phagocytic capacity is associated with the M2-like polarization state (50).To determine whether secreted signals by compressed macrophages influenced uncompressed cells, we treated macrophages grown in standard culture conditions with conditioned media from compressed or uncompressed macrophages. We found a significant decrease in phagocytosis by macrophages treated with compressed macrophage media, indicating potentially competing primary and secondary effects of compression (**Figure 6C**). Secreted nitric oxide (NOS), a mediator of macrophage cytotoxic activity, was measured via a Griess assay of compressed macrophage-conditioned media. After 48 hours of compression, there was a statistically significant increase in the nitrite, a derivative of secreted NOS, in conditioned media (**Figure 6D**). While NOS production is a canonically M1-like marker, its secretion by TAMs may promote treatment resistance (51).

**Figure 6.**
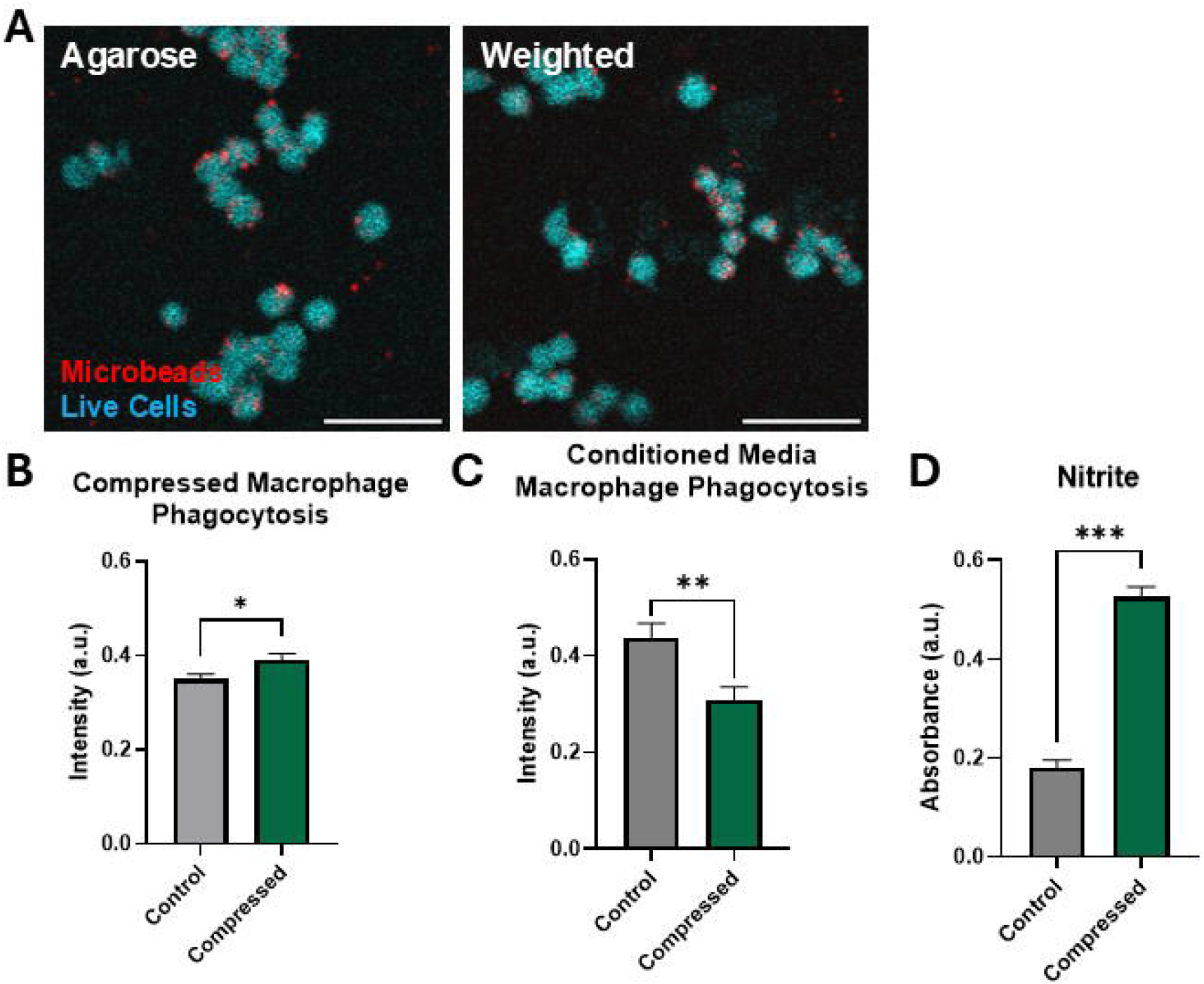
Macrophages under compression have increased phagocytic capacity and nitric oixide production. (**A**) Representative images of macrophages (cyan) and fluorescent microbeads (red). After 24 hours of compression, macrophages were incubated with fluorescent microbeads, internalizing them based on phagocytic capacity. Scale bar represents 50 µm. (**B**) Quantification of intensity of intracellular fluorescent microbeads phagocytosed after compression as a metric of the overall phagocytic capacity (N=12-14 images per group). (**C**) Quantification of macrophage microbead uptake when cultured with conditioned media from compressed macrophages (N=9 images per group). (**D**) Quantification of nitrite (indicative of NOS) released by macrophages after 48h of compression (N=3 samples per group). *p ≤ 0.05, **p ≤ 0.01, ***p ≤ 0.001.

## Discussion

Here, we establish a clear relationship between the mechanical TME and its immune residents. The direct contribution of macrophages to tumor progression, immune escape, and patient outcome has been extensively studied and is well-reviewed elsewhere (4,52–54). Macrophages are known to respond to their physical environment, including matrix stiffness and topography, cyclic tension or compression, and shear stress (55,56). However, studies on compressive solid stress on macrophages have generally been limited to the context of bone or orthodontic intervention, so the magnitudes of solid stress and the environments in which they are studied differ drastically from those found in the TME (16,17,57). *Ex vivo* measurements of murine tumors reveal a range of 0.1 to 10 kPa maximum solid stress, with murine GBMs falling in the range of 0.1 to 0.2 kPa (11,30,31). We therefore designed our custom 3D-printed weight to apply 0.14 kPa of solid stress uniformly across its surface. This study is the first of its kind to study chronic compressive stress on macrophages in a custom compression system designed to mimic the mechanical stress in the GBM TME.

For nearly every metric we applied, compression caused alterations resembling both M1-like and M2-like polarization states (**Figure 7**). For example, increased macrophage size, seen under compression, is a typical characteristic of M1-like macrophages (32). However, the more irregular cell shape (characterized by circularity and solidity) is more commonly associated with the M2-like phenotype (32). Polarization of macrophages by added cytokines has been observed to alter macrophage elongation, but compressed macrophages had no significant change in elongation or eccentricity (data not shown) (58). Macrophage morphology is not only an indicator of polarization but can also directly impact it as part of a two-way interaction. For example, macrophages forced to adopt an elongated shape altered macrophage polarization and sensitivity to pro-inflammatory stimuli relationship (58). In our system, the unconfined, uniaxial compression may have physically forced the cells to spread thinner, resulting in the observed increased area. This forced change in shape may itself be one mechanism by which compression drives changes in macrophage phenotype.

**Figure 7.**
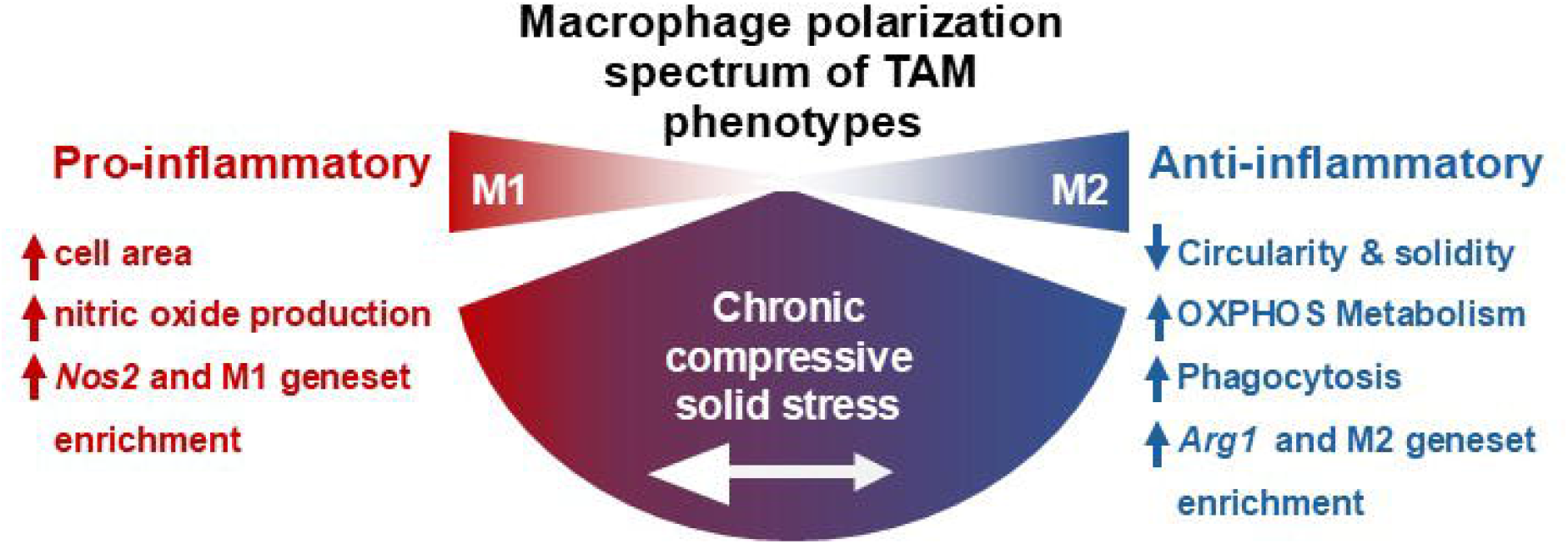
Overview of the effect of compression on macrophages. Compression causes macrophages to adopt both pro- and anti-inflammatory associated traits, with the net effect of promoting an immunosuppressive, tumor-associated and glioma-associated macrophage phenotype.

As expected, compression was accompanied by a clear transcriptional response. Based on gene ontology analysis, the biological and molecular processes upregulated under compression were largely related to chemotaxis and cytokine signaling, while many of the most downregulated processes were involved in cell replication. Visualizing these two processes in a protein-protein interaction map highlighted IL-1β as one of the central hubs of interaction for both. IL-1β participates in a feedback loop in several human cancers, promoting tumor immune escape via increased PD-L1 expression, the secretion of macrophage-recruiting proteins, the acquisition of malignant characteristics by tumor cells, and worse patient prognosis (59–62). In GBM in particular, macrophage-derived IL-1β promotes edema, induces GBM tumor growth, promotes cancer stem cell phenotype, and leads to further immune reprogramming by GBM cells including macrophage polarization and recruitment (63–67). The induction of IL-1β production by macrophages is attributed to factors secreted by tumor cells, but here we show that compression alone, in the absence of any tumor-derived biochemical signals, causes an increase in *Il1b* gene expression. IL-1β may therefore be an important intermediary between the pathological mechanical microenvironment and macrophage immune reprogramming.

We note that compression caused a simultaneous decrease in pathways associated with cell cycling and proliferation, and an increase in genes associated with apoptosis. Together, these would suggest that compression may induce a decrease in the macrophage population in a closed system. Weak macrophage proliferation within tumors is one of the biggest hurdles for engineered macrophage-based therapies, such as CAR-macrophages (68). However, the concurrent increase in pathways related to migration and chemotaxis suggests that an open system, such as a tumor *in vivo*, may still accumulate macrophages in response to compression via recruitment from the bloodstream and migration into the tumor. Compressed macrophages are not only potentially more migratory themselves, but their upregulation of monocyte chemoattractants such as CCL2 and CCL5 (constituents of the migration-associated geneset) indicate that they could play an active role in the direct recruitment of more macrophages from the bloodstream, as TAMs are observed to do in solid tumors (69).

Macrophage polarization did not neatly align with either the M1 or M2 side of the canonical polarization spectrum. As has been observed in gliomas, compressed macrophages upregulated both M1-like and M2-like genes (70). Compression did, however, consistently cause upregulation of TAM and GAM genesets, except for the first pro-inflammatory TAM geneset. Patients with tumors enriched in this subtype responded better to immunotherapy than those enriched in the pro-tumor subtype, whereas the patients with macrophages matching the pro-tumor geneset (as compressed macrophages do) had worse outcomes (71). Functionally, compressed macrophages were more phagocytic, in line with gene expression data, and released more nitric oxide into their media. Future work involving co-culture with glioma cells would help determine whether this corresponds to a change in anti-tumor activity. Elements of the IL6/JAK/STAT3 signal pathway were also upregulated, which are associated with worse prognosis in solid tumors and important in regulating macrophage response to stimuli (72,73). Compressed macrophages also upregulated ECM remodeling-associated genes, including Lox and several MMPs. Among these is MMP9, also a prominent hub protein in our protein-protein interaction map. MMP9 promotes cancer cell invasion and angiogenesis and is associated with poor patient prognosis in solid tumors (74).

The metabolic state of compressed macrophages was more difficult to determine. M2-like macrophages have been observed to have a longer average fluorescence lifetime than M1-like macrophages, a characteristic that can help distinguish the two with relatively high accuracy (49,75). The increased fluorescence lifetime of compressed macrophages thus suggests a shift towards a more M2-like metabolic state. The increase in fluorescence lifetime was expected to correspond with more reliance on oxidative phosphorylation, and less on glycolysis (76). As noted by others, average fluorescence lifetime alone is influenced by factors other than oxidative or glycolytic cell state, so more comprehensive FLIM analysis or other metabolic assays would be required to more fully understand cellular metabolic state (77). Interestingly, hypoxic conditions are expected to be accompanied by a decrease in cell fluorescence lifetime (78,79), but compressed macrophages simultaneously upregulated hypoxia genes and had increased average fluorescence lifetime. Macrophages polarized to an M1-like state by interferon-gamma relies on increased glycolysis, which allows for Hif-1a, IL-1b, and nitric oxide production and JAK/STAT signaling - all consistent with our gene expression and media analysis (80). Together, these results highlight the complexity of macrophage polarization and metabolism, and highlight the need for further mechanistic studies linking the two in the context of chronic compression.

Targeting myeloid cells has shown great promise as part of a combinatorial approach, enhancing the effect of other strategies to alter the TME. For example, macrophages are essential for the full beneficial effect of dual inhibition of Ang-2 and VEGF in murine GBM (81). Targeting myeloid cells in the GBM TME by CCR2 inhibition is sufficient to remove the barrier to ICB efficacy and extends survival in murine GBM models (82). Reprogrammed macrophages may also serve a as powerful therapeutic tool, as pharmacologically normalizing the tumor vasculature results in improved survival mediated by reprogrammed tumor-resident macrophages (83). This strategy is advantageous because it can target myeloid cells in the bone marrow and therefore does not need to cross the BBB to have effect. Targeting solids tress in the TME will also interrupt the immune-mechanical feedback loop. Reducing any of the solid components of the tumor (tumor cells, stromal cells, or ECM) causes a reduction in solid stress (84). Losartan, an angiotensin system inhibitor, can help control ICB-related adverse events its use is associated with improved overall survival in both murine and human GBM (85,86). This may be due to its inhibition of collagen/hyaluronan synthesis, CCR2 signaling, and monocyte recruitment, potentially simultaneously impacting both TAMs and the mechanical TME (87,88). Future studies on the effect of chronic compression *in vivo* using a compressive cranial window will elucidate the more complex interaction between mechanical force, myeloid and other immune cells, and the brain microenvironment (89).

In conclusion, we have demonstrated that chronic mechanical compression alone can drive macrophages towards a complex phenotype largely resembling that seen in GBM and other cancers. We previously established that macrophages themselves can contribute to the generation of solid stress (15). This work provides evidence to close the positive immunomechanical feedback loop between macrophages and solid stress, wherein macrophages may both generate and respond to solid stress in a manner that contributes to immune suppression and tumor progression. This cycle represents a promising therapeutic target, as reducing solid stress, macrophage sensing and response to it, or macrophage contribution to it, could break the cycle and normalize the tumor immune microenvironment.

## Acknowledgements

Funding for this work was provided by the National Institutes of Health (NIH/ NIGMS R35GM151041 to M.D.), and the Walther Cancer Foundation Interdisciplinary Interface Training Project (IITP) Grant and the Edison Innovation Fellowship (to A.A.B.). RNA sequencing was done through the Notre Dame Genomics and Bioinformatics Core Facility. The authors thank Ms. R’nld Rumbach for technical assistance, and Dr. Donny Hanjaya-Putra and his graduate students Mr. Donghyun Paul Jeong and Ms. Eva Hall for imaging assistance. We thank Maksym Zarodniuk for advice regarding RNA extraction and analysis.

## Data availability statement

RNA sequencing raw data will be deposited in a public repository before final publication.

## Author Contributions

AB: Conceptualization, Investigation, Data curation, Methodology, Formal analysis, Funding acquisition, Visualization, Writing – original draft, review & editing. HC: Investigation, Data curation, Formal Analysis, Visualization, Writing – review & editing. JN: Formal Analysis, Visualization, Writing – review & editing. SH: Conceptualization, Methodology, Supervision, Resources, Writing - review & editing. MD: Conceptualization, Methodology, Supervision, Project administration, Funding acquisition, Resources, Writing – original draft, review & editing.

